# APOLLO: A genome-scale metabolic reconstruction resource of 247,092 diverse human microbes spanning multiple continents, age groups, and body sites

**DOI:** 10.1101/2023.10.02.560573

**Authors:** Almut Heinken, Timothy Otto Hulshof, Bram Nap, Filippo Martinelli, Arianna Basile, Amy O’Brolchain, Neil Francis O’Sullivan, Celine Gallagher, Eimer Magee, Francesca McDonagh, Ian Lalor, Maeve Bergin, Phoebe Evans, Rachel Daly, Ronan Farrell, Rose Marie Delaney, Saoirse Hill, Saoirse Roisin McAuliffe, Trevor Kilgannon, Ronan M.T. Fleming, Cyrille C. Thinnes, Ines Thiele

## Abstract

Computational modelling of microbiome metabolism has proved instrumental to catalyse our understanding of diet-host-microbiome-disease interactions through the interrogation of mechanistic, strain- and molecule-resolved metabolic models. We present APOLLO, a resource of 247,092 human microbial genome-scale metabolic reconstructions spanning 19 phyla and accounting for microbial genomes from 34 countries, all age groups, and five body sites. We explored the metabolic potential of the reconstructed strains and developed a machine learning classifier able to predict with high accuracy the taxonomic strain assignments. We also built 14,451 sample-specific microbial community models, which could be stratified by body site, age, and disease states. Finally, we predicted faecal metabolites enriched or depleted in gut microbiomes of people with Crohn’s disease, Parkinson disease, and undernourished children. APOLLO is compatible with the human whole-body models, and thus, provide unprecedented opportunities for systems-level modelling of personalised host-microbiome co-metabolism. APOLLO will be freely available under https://www.vmh.life/.

## Introduction

The human microbiome plays an important role in health and wellbeing (Lynch and Pedersen, 2016). Changes in gut microbiome composition have been associated with numerous diseases, including cardiometabolic, gastrointestinal, and neurological (Hou et al., 2022; Lavelle and Sokol, 2020). Other human microbiomes, such as the skin and vaginal microbiomes, have received relatively less attention but have also been linked to diseases, such as acne vulgaris, atopic dermatitis, bacterial vaginitis, and polycystic ovarian syndrome (Byrd et al., 2018; Han et al., 2021). The microbiome composition varies both by body site and individual (Ruan et al., 2020) and is influenced by a multitude of factors, including genetics, sex, diet, lifestyle, geography, and ethnicity (Alexander and Turnbaugh, 2020; Rothschild et al., 2018). Particularly, a “Westernised” lifestyle characterised by an industrialised environment, a sugar- and fat-rich diet, sanitation, lack of exercise, and antibiotics overuse has been associated with detrimental changes in microbiome composition, such as a loss in taxa performing functions considered beneficial to the host, e.g., butyrate production (Sonnenburg and Sonnenburg, 2019). Studying the microbiomes of non-industrialised populations may help to analyse the influence of the Westernised lifestyle on the microbiome (Smits et al., 2017; Sonnenburg and Sonnenburg, 2019). Moreover, individuals from the Global South are still understudied, limiting insights into the specific role of their microbiomes in health and disease states, as most sequencing efforts have focussed on populations from North America and Europe (Tamburini et al., 2022).

Advances in method development have enabled the retrieval of metagenome-assembled genomes (MAGs) directly from metagenomic sequencing data (Kang et al., 2015; Quince et al., 2017). Recent studies have assembled MAGs from thousands of human microbiome samples, resulting in tens of thousands of strain-resolved microbial genomes, including many yet uncultured and uncharacterised organisms (Almeida et al., 2019; Pasolli et al., 2019). Pasolli et al. reconstructed over 150,000 MAGs from 9,428 publicly available metagenomes spanning 32 countries, five body sites, and all age groups (Pasolli et al., 2019). In another study, Almeida et al. reconstructed over 90,000 MAGs from 11,850 human gut microbiome metagenomes spanning six continents (Almeida et al., 2019). These resources provide an unprecedented opportunity to elucidate the metabolic capabilities of currently still understudied taxa as well as the influence of factors, such as age, geography, and lifestyle, on the structure and functions of the human microbiome. Investigating the relationship between human microbiomes and host traits, such as age, diet, lifestyle, health status, and ethnicity, usually requires multivariate statistics, machine learning methods as well as computational systems biology approaches (Heinken et al., 2021a; Sudhakar et al., 2021).

Constraint-based reconstruction and analysis (COBRA) enables mechanistic, strain-, and molecule-resolved metabolic modelling by integrating meta-omics data (Heirendt et al., 2019). COBRA relies on high fidelity genome-scale metabolic reconstructions of the organism(s) of interest (Palsson, 2015). These genome-scale reconstructions can be converted into mathematical models through the application of condition-specific constraints (e.g., a given nutrient environment), and then interrogated in simulations (Palsson, 2015). By joining microbial reconstructions into a sample-specific microbial community model and predicting the microbiome-wide metabolic fluxes, the metabolic potential of a given microbiome can be elucidated *in silico* (Heinken et al., 2021b). Importantly, since COBRA is based on molecule-resolved metabolic networks, it can mechanistically predict links between microbial enzymes and faecal or blood metabolites (rather than through statistical associations) (Hertel et al., 2021).

We have previously developed the AGORA resources of human gut microbial reconstructions (Heinken et al., 2023; Magnusdottir et al., 2017). AGORA2 accounts for 7,302 strains from 25 phyla and accurately captures the distribution of metabolic pathways across members of the human gut microbiome (Heinken et al., 2023). Thus far, AGORA reconstructions have enabled over 50 studies interrogating microbe-microbe, host-microbe, and microbiome interactions (Heinken et al., 2021b). For the generation of the AGORA2 resource, a semi-automated curation pipeline, DEMETER (Heinken et al., 2021d), has been developed. DEMETER takes draft reconstructions, e.g., from KBase (Arkin et al., 2018) or ModelSEED (Faria et al., 2018), and converts them into the virtual metabolic human (VMH) name space (Noronha et al., 2019). These draft reconstructions are then subjected to a set of quality control tests, including testing for flux and stoichiometric consistency, for mass-and charge-balance, for having correct reconstruction structure, and for producing realistic amounts of biomass and ATP (Heinken et al., 2023; Lieven et al., 2020; Thiele and Palsson, 2010b). Furthermore, the draft reconstructions can be gap filled based on refined genome annotations and experimental data from >600 studies and books for more than 1,000 species (Heinken et al., 2021d). DEMETER generated reconstructions perform better when compared to independent experimental data than reconstructions generated by other available (semi-) automated reconstruction tools (Agren et al., 2013; Heinken et al., 2023; Machado et al., 2018; Zimmermann et al., 2021). Hence, DEMETER is an attractive choice for scaling the reconstruction and analysis to an even larger resource of human microbial genomes.

## Results

We present APOLLO, a microbial reconstruction and analysis resource encompassing 247,092 genome-scale metabolic reconstructions built from two separate MAG resources (Figure 1). We systematically interrogated each strain-level reconstruction using constraint-based modelling. Subsequently, we developed a machine learning classifier to predict taxonomic strain assignment based on model properties., i.e., reaction presence and absence and metabolite production capabilities. Finally, we built 14,451 personalised microbiome models and interrogated them through simulations. Machine learning and statistical analyses identified key features, which could stratify individual microbiomes based on diseases, body site, and age group. Taken together, APOLLO currently represents the largest resource of genome-scale reconstructions on strain- and microbiome-level to date. Furthermore, we illustrate examples for the many potential applications of the APOLLO resource to systematically investigate metabolic capabilities on strain and microbiome level.

**Figure 1:**
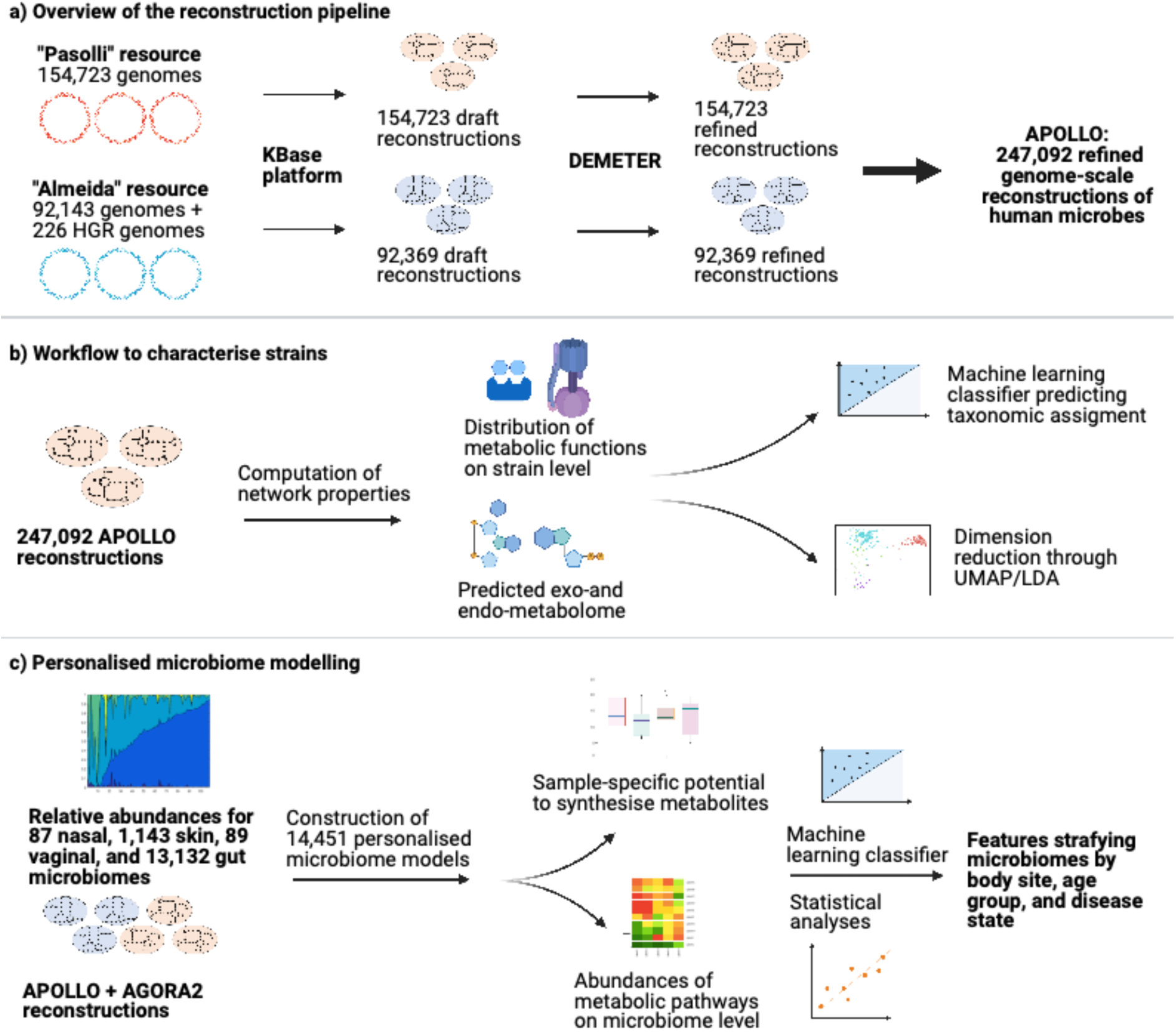
Overview of the reconstruction and analysis pipeline to construct and interrogate APOLLO. a) Overview of the pipeline to reconstruct the “Pasolli” and “Almeida” MAG resources, consisting of generation of draft reconstructions through KBase, and refinement, testing, and debugging of draft reconstructions through DEMETER. b) Overview of the workflow to systematically characterise the distribution of features across taxa. For all APOLLO strains, metabolic functions were systematically computed. Through a machine learning classifier, taxonomic assignment of strains was predicted based on the computed features. Strains were clustered based on metabolic similarities using UMAP/LDA. c) Creation and interrogation of personalised microbiome models. Mapped relative abundances were used to construct models for 14,451 microbiomes from four body sites. Reaction abundance and presence on microbiome level were determined, and metabolite production potential was computed for a subset of samples. Key differences between groups were identified through machine learning and statistical analyses. Created with BioRender.com.

### APOLLO: a resource of 247,092 refined genome-scale reconstructions

To build genome-scale metabolic reconstructions from the two MAG resources, we retrieved 154,723 MAGs (Pasolli et al., 2019) (Pasolli resource, Table S1) and 92,143 MAGs (Almeida et al., 2019) (Almeida resource, Table S1), as well as 226 genomes from the Human Gastrointestinal Bacteria Culture Collection (Forster et al., 2019) that served as an independent validation dataset (Table S1). For each genome, a draft reconstruction was generated via the KBase online platform (Arkin et al., 2018) and subsequently, refined in a semi-automated manner using the DEMETER pipeline (Heinken et al., 2021d) (Figure 1a, Methods). Where possible (57.0% and 45.91% of reconstructions for Pasolli and Almeida, respectively, representing 52.85% of reconstructions in total), the metabolic reconstructions were expanded and refined based on available experimental data in the scientific literature for over 1,000 species (Methods). Through the DEMETER test suite (Heinken et al., 2021d), we ensured that the refined reconstructions conformed with established standards in the constraint-based modelling field (Lieven et al., 2020; Thiele and Palsson, 2010b), and, where possible, captured the known traits of the target organisms, including the presence of the periplasm, where appropriate. Altogether, we generated and tested 247,092 semi-automatically refined genome-scale reconstructions, deemed APOLLO, which are available at the Virtual Metabolic Human (https://www.vmh.life/) (Noronha et al., 2019).

### High-quality genome-scale reconstructions derived from MAGs

Most genome-scale reconstructions to date have relied on medium-to high-quality reference genomes, with only few studies using MAGs as the starting point for reconstruction (Frioux et al., 2020; Zorrilla et al., 2021). To assess the predictive potential and quality of MAG-derived genome-scale reconstructions, we compared their features between the Pasolli- and Almeida-derived reconstructions with the 7,302 high-quality AGORA2 reconstructions (Heinken et al., 2023). AGORA2 has been built from reference genomes with an additional step involving refinement of genome annotations based on manually performed comparative genomic analyses (Heinken et al., 2023). On average, the refined Pasolli and Almeida reconstructions contained 997.92 (± 215.4) reactions, 955.19 (±161.81) metabolites, and 534.13 (±170.86) genes (Figure 2a, b), thus, being smaller than the AGORA2 reconstructions. Combined, they also had a lower number of unique reactions and metabolites (Figure 2a, b). Both features are a direct consequence of using MAGs, which are typically incomplete without further curation (Chen et al., 2020) rather than reference genomes, for the generation of the draft reconstructions. Additionally, the lack of comparative genomic analyses, particularly for drug metabolism-specific pathways, which accounted for over 2,000 unique reactions in AGORA2 (Heinken et al., 2023), resulted in a smaller number of unique reactions in APOLLO (Figure S1). As many genomes in the Pasolli and Almeida resources could not be taxonomically assigned to a species, but only to the genus level or higher (Almeida et al., 2019; Pasolli et al., 2019), less experimental data was overall available for the refinement of the reconstructed strains compared to AGORA2. We determined the strains agreeing with all available experimental data used for refinement (Methods) for draft and refined reconstructions of Pasolli and Almeida strains. In total, 81.03% and 73.14% of refined Pasolli and Almeida reconstructions, respectively, agreed with the available experimental data, compared with 11.30% and 9.65%, respectively, of the draft reconstructions (Table S2). This result demonstrates an improvement in predictive potential through the performed refinement. We predicted the maximal possible growth rates and ATP yields on aerobic and anaerobic complex medium, which were within biologically reasonable ranges for all reconstructions (Figure 2a). The average percentage of flux-consistent reactions, i.e., reactions that can carry a non-zero flux in the simulation conditions, was higher in refined APOLLO reconstructions compared to draft reconstructions (63.18% +/- 6.18 vs. 60.93% +/-4.17), but lower than in AGORA2 reconstructions (67,77% +/- 7.37, Figure 2a, Table S3). Taken together, we found that the reconstructions in the APOLLO resource generated from two separate, independent MAG collections were comparable in their properties and overall characteristics. Compared with AGORA2, the metabolic coverage and percentage of flux-consistent reactions were smaller, but the quality of the reconstructions in terms of thermodynamic feasibility and realistic growth rates was similar. The detailed characteristics for each APOLLO reconstruction are shown in Table S3.

**Figure 2:**
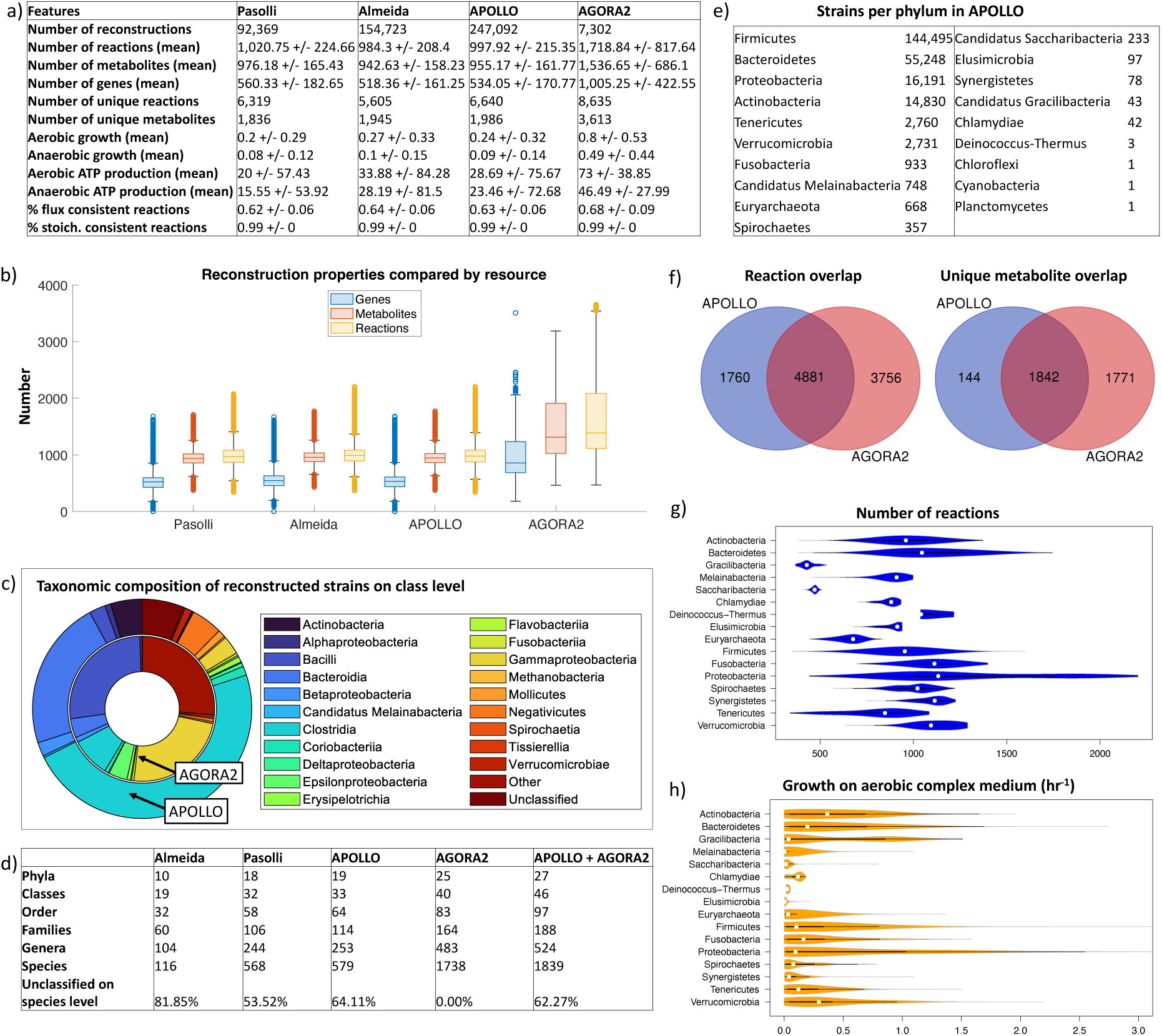
Overview of characteristics of APOLLO. a) Comparison of reconstruction features between Pasolli reconstructions, Almeida reconstructions, APOLLO, and AGORA2. b) Numbers of reactions, metabolites, and genes across Pasolli reconstructions, Almeida reconstructions, APOLLO, and AGORA2. c) Taxonomic assignment in APOLLO and AGORA2 plotted as fractions of total contained strains. d) Number of unique taxa from phylum to species and fraction of unclassified strains on the species level in Pasolli reconstructions, Almeida reconstructions, APOLLO, and AGORA2. e) Number of strains in phyla in APOLLO, rank ordered. f) Overlap in reaction and unique metabolite content between all APOLLO and AGORA2 reconstructions. g-h) Taxon-specific computed features for strains contained in APOLLO. g) Number of reactions, h) predicted growth rate (hr^-1^) on aerobic complex medium.

### Taxonomic coverage and intraspecies metabolic variability in APOLLO

Next, we examined the taxonomic coverage of the 247,092 APOLLO metabolic reconstructions. Overall, the taxonomic distribution was close to a typical human microbiome composition (Lloyd-Price et al., 2016), with Bacteroidia and Clostridia classes accounting for most strains (Figure 2c). However, APOLLO contained fewer unique taxa than AGORA2 despite a much higher number of strains (Figure 2d), likely due to the high number of unclassified strains. Moreover, the selection of reconstructed genomes in AGORA2 had been done manually in a literature- and data-driven manner resulting in a high number of unique species (Heinken et al., 2023). The Pasolli resource had a greater overlap with AGORA2 and a higher number of unique taxa than the Almeida resource (Figure S2). As expected, Firmicutes, Bacteroidetes, Actinobacteria, and Proteobacteria were the most abundant of the 19 phyla with a total of 230,764/247,092 strains (93.39%, Figure 2e). Of the 6,641 unique reactions and 1,986 unique metabolites contained within APOLLO, the majority were shared with AGORA2 (Figure 2f). Moreover, the overall content of reconstructions generated from the Almeida and Pasolli MAG resources were comparable to each other (Figure S2). AGORA2 contains 3,756 reactions and 1,771 metabolites not found in APOLLO (Figure 2e), the majority of which belonged to peripheral pathways, such as xenobiotics and polyphenol metabolism (Figure S3). These pathways, absent in APOLLO, were due to AGORA2 stemming from reference genomes and its additional curation through comparative genomics analyses. As already observed for AGORA2 (Heinken et al., 2023), reaction numbers and achieved growth rates on a complex medium greatly varied between phyla (Figure 2g-h). To estimate to which extent reactions were shared between strains, we determined the pan-and core-reactome for each species, i.e., the reactions found in at least one strain and in all strains of the species, respectively (Table S4). For the 30 species with the highest number of strains in APOLLO, the pan-reactomes varied from 1,807 to 3,092 reactions and only 42.39% to 63.07% of these pan-reactome reactions were contained in the respective core reactomes (Figure S4), indicating high intraspecies metabolic variety. This result is in line with a previous analysis of MAGs reporting that 38.5–57.6% of a species’ pan-genome was included in its core genome (Zorrilla et al., 2021).

### Metabolic features are predictive of taxonomic assignment of the reconstructed strains

Beyond reaction and pathway presence, to explore the functional metabolic potential, we predicted the uptake and secretion potential (representing the *in silico* exometabolome), and the intracellular metabolite production potential (respresenting the *in silico* endometabolome) of each APOLLO reconstruction using flux balance analysis (Orth et al., 2010), assuming unlimited uptake of nutrients (Methods). In total, 517 could be consumed and 434 secreted (of which 360 overlapped), and 1,248 could be synthesised intracellularly by at least one strain (Table S5). To reduce the dimensionality of the APOLLO dataset, we visualised the separation of strains based on reaction presence, metabolite uptake and secretion potential, and internal metabolite production potential in 2D via linear discriminant analysis (LDA), and in 3D via uniform manifold approximation and projection for dimension reduction (UMAP) (McInnes et al., 2018). For both reaction content and predicted metabolic capabilities, the two methods demonstrated a clear separation of strains based on their taxonomic assignment to phyla (Figures 3a-c, S5-S7) as well as on lower taxonomical levels (Figures S8-S10). Separation into clusters by taxon was also observed for the Pasolli and Almeida reconstructions separately (Figures S11-S12). The previously demonstrated (Cabbia et al., 2020; Heinken et al., 2023; Magnusdottir et al., 2017) taxonomic relatedness of strains based on similarity in reaction content and predicted metabolic capabilities was also a feature of the much larger APOLLO resource.

**Figure 3:**
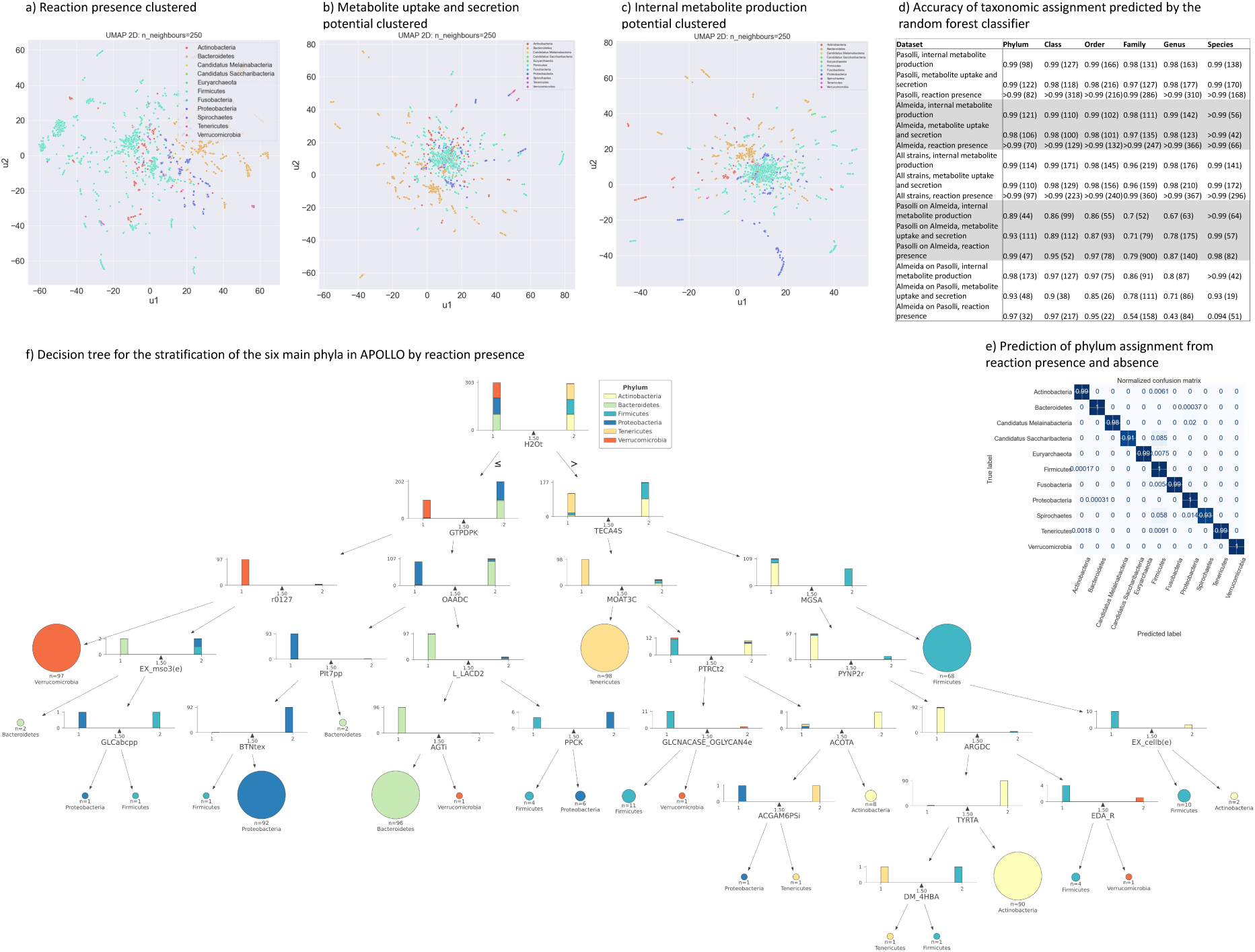
Analysis of strain-level reconstruction features in APOLLO through dimension reduction and the random forests classifier. a-c) Clustering of strain-level predicted model properties through UMAP on the class level. a) Reaction presence, b) metabolite uptake and secretion potential, c) internal metabolite production potential. UMAP analyses on order level for the same data are shown in Figure S8-10. d) Overview of taxonomic assignment for the three datasets and from phylum to species predicted by the random forest classifier. Shown is the accuracy of the predicted taxonomic assignment against the assignment reported by the original authors (where classification was possible). The number of features that were sufficient to achieve the reported accuracy is shown in brackets. The list of classifying features for each prediction is available in Table S6. e) Normalised confusion matrix for the prediction of phylum assignment based on reaction presence and absence for phyla with more than 125 representatives in APOLLO. f) Decision tree showing the minimal set of reactions leading to strain classification by phylum by their presence or absence. Shown are results generated from 100 randomly selected strains for each phylum. 1 = Absence of the reaction, 2 = presence of the reaction. Reactions are shown by VMH ID (https://www.vmh.life) (Noronha et al., 2019).

To test whether the computed properties and metabolic capabilities of the models could be predictive of the reconstructed genomes’ taxonomic assignments, we developed a machine learning classifier (Methods). The classifier was applied to all computed datasets (i.e., reaction presence, uptake and secretion potential, and internal metabolite production potential), on multiple taxon levels (i.e., species, genus, family, order, class, and phylum), and on the following scenarios: (i) Pasolli reconstructions only, (ii) Almeida reconstructions only, (iii) Almeida reconstructions using Pasolli reconstructions as the training set, (iv) Pasolli reconstructions using the Almeida reconstructions as the training set, and (v) all APOLLO reconstructions. Classifiers were generated for taxa encompassing at least 125 strains, with a split of 80:20 for training set and validation set (Methods). In most of the considered cases, the generated classifiers were highly predictive on all taxonomic levels and for both reaction presence and predicted metabolic capabilities (Figure 3d). For instance, the presence and absence of only 82 out of the 6,319 unique reactions across the Pasolli reconstructions predicted taxonomic assignment on phylum level for 11 phyla with >99% accuracy (Figure 3d). In fact, all taxonomical levels for the three datasets were predicted with at least 97% accuracy for Pasolli reconstructions (Figure 3d). Similar predictive potential was achieved for Almeida reconstructions, and for the complete APOLLO resource (Figure 3d). When the Pasolli reconstruction datasets were used as the training set to develop the classifier, the taxonomy of Almeida reconstructions could be predicted with an accuracy of 86% or higher on most taxonomical levels with only family and genus performing more poorly (Figure 3d). Similarly high performances were achieved on most taxonomical levels and datasets when using the Almeida reconstruction data as the training set on the Pasolli reconstruction data (Figure 3d). Taken together, using a machine learning classifier we demonstrated that, in addition to reconstructions clustering together by taxonomic similarity based on their reaction presence and predicted metabolic capabilities, the same model-specific features were highly predictive of taxonomic assignment on all levels (species to phylum). The Pasolli reconstruction datasets could serve as a training set for the Almeida reconstruction and vice versa to an extent highlighting that the content of both reconstruction resources was overall comparable despite the different sources of input MAGs.

For the entire APOLLO resource, 97 out of 6,640 unique reactions could predict the phylum assignment for 11 phyla with >99% accuracy (Figure 3d, Table S6). The confusion matrices demonstrated that four phyla were predicted by the presence and absence of these reactions with 100% accuracy and further four with 99% accuracy (Figure 3e). Similarly, the metabolite uptake and secretion potential, and the internal metabolite production potential was highly predictive of assignment for most phyla (Figure S13-S14). To gain insight into the reactions that are indicative of phylum assignment, we generated a decision tree based on presence and absence of reactions using 100 randomly chosen reconstructions from the five most abundant phyla (Figure 3f). For instance, the absence or presence of the H_2_O transporter via diffusion from extracellular environment to cytosol (VMH ID: H2Ot) assigned strains to Actinobacteria, Tenericutes, and the majority of Firmicutes on one side, representing gram-positive taxa, and Bacteroidetes, a subset of Firmicutes, Proteobacteria, and Verrucomicrobia on the other side, representing gram-negative taxa (Figure 3f). Furthermore, nearly all Verrucomicrobia representatives (97/100) split from other gram-negative phyla based on the absence of two reactions, GTP diphosphokinase and glutaminase-asparaginase (VMH IDs: GTPDPK, r0127). Most Proteobacteria representatives (92/100) could be distinguished from Bacteroidetes representatives by the absence of two reactions, namely oxaloacetate decarboxylase and a periplasmatic phosphate symporter (VMH IDs: OAADC, PIt7pp), and the presence of a periplasmatic biotin transporter (VMH ID: BTNtex). These examples demonstrate that a small number of reactions were sufficient to correctly assign the taxon-level in a subset of strains and highlight taxa-specific metabolic differences are captured in the APOLLO resource.

### Generation of 14,451 personalised microbiome models varying in body site, age group, and host health status

Through the integration of metagenomics data, genome-scale metabolic models can give rise to personalised community models of microbiomes (Heinken et al., 2021a). To analyse microbiome-wide structure and functions of the microbiome with respect to age group, health status, and country of origin of the corresponding human hosts, we built 13,332 gut microbiome community models encompassing all age groups, 21 countries, and 21 diseases (Methods, Table S7). Moreover, to delineate microbiome structure and function in distinct body sites, we built 87 nasal cavity, 1,143 skin, and 89 vaginal microbiome community models (Methods, Table S7). The resulting 14,451 personalised microbiome models contained on average 158,007±60,803 reactions, 148,575±59,715 metabolites, and 143.89±65.10 mapped strains (Table S7). We used these personalised microbiome models to investigate their emerging metabolic traits.

### The machine learning classifier identifies features that best stratify microbiomes

To address the question whether there exists any metabolic trait that can characterise the different microbiome samples, we defined 11 datasets consisting of two to ten stratification groups, including eight diseases, four body sites, ten countries of origin, and two age groups (Figure 4a, Table S8, Methods). We determined the microbiome-level absolute presence and relative abundance of each reaction as well as the relative abundance of subsystems (Methods, Table S9a-c). We then generated machine learning classifiers to predict group assignments based on relative strain-level abundance, relative reaction abundance, absolute reaction presence, and relative subsystem abundance (Methods, Figure 4a, Table S10). The nasal cavity, skin, and vaginal microbiomes could be distinguished based on the metabolic model features with 99% accuracy and 97% based on strain-level abundance (Figure 4a). Inflammatory bowel disease (IBD) and healthy control gut microbiomes could be stratified with 99% accuracy based on two to nine signature features (Figure 4a). Similarly, the gut microbiomes of Parkinson disease (PD) patients and healthy controls could be distinguished with 92% accuracy (Figure 4a). Healthy and premature infant gut microbiomes, and gut microbiomes of healthy adults and adults with infection were also well-predicted with 98% accuracy. Group assignments in other datasets were less accurately predicted (Figure 4a). A clear visual separation into distinct clusters was observed for nasal cavity, skin, and vaginal microbiomes of healthy adults, gut microbiomes of healthy adults and infants, gut microbiomes of healthy and premature infants, gut microbiomes of healthy and undernourished infants, gut microbiomes of healthy adults and adults with infection, and PD and healthy control gut microbiomes (Figure 4b-g). Less visual separation was observed for gut microbiomes of healthy individuals by country, of patients with infection with and without antibiotics use, of normal weight and obese individuals, and of individuals with and without type 2 diabetes (T2D) (Figures S15-18), in agreement with the observed lower predictive potential of the classifiers for these datasets (Figure 4a).

**Figure 4:**
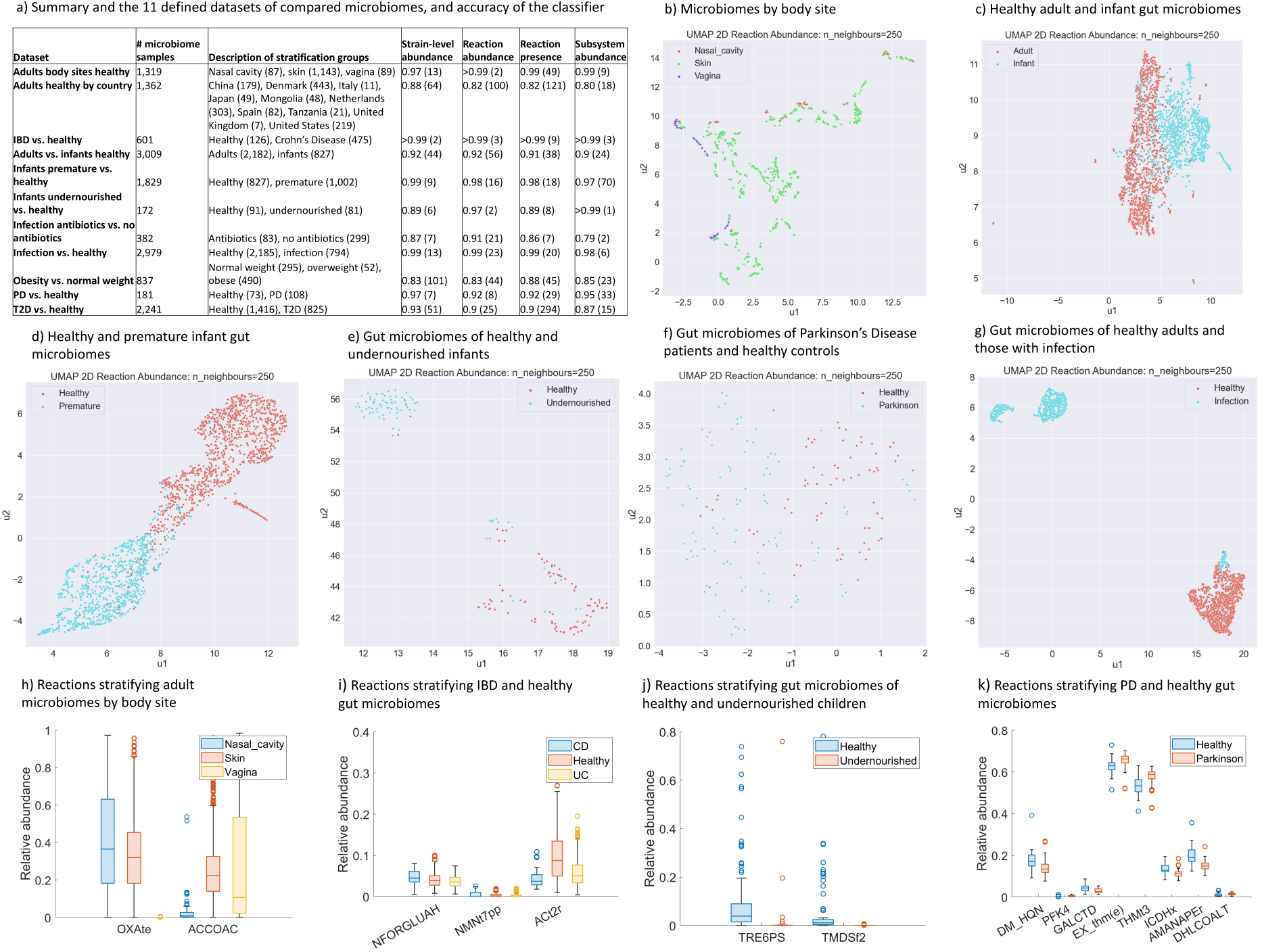
Analysis of personalised microbiome models constructed from APOLLO through dimension reduction and the random forests classifier. a) Overview of the 11 modelled microbiome datasets and accuracies obtained through random forests analyses. For the sample inclusion criteria, see Methods. Shown are the accuracies for the correct prediction of stratification groups by the random forests classifier for each dataset based on strain-level relative abundance as well as reaction abundance, reaction presence, and subsystem abundance summarised for each sample-specific model. The number of features that were sufficient to achieve the reported accuracy is shown in brackets. The list of classifying features for each prediction is shown in Table S10. b-g) Clustering of microbiome model datasets defined in this study by body site through UMAP by relative reaction abundance. b) Healthy microbiomes by body site, c) healthy adult and infant gut microbiomes, d) healthy and premature infant gut microbiomes, e) gut microbiomes of healthy and undernourished children, f) gut microbiomes of PD patients and healthy controls, g) gut microbiomes of healthy adults and those with infection. h-k) Subsets of reactions that predicted stratification group by random forests analysis. Shown are the VMH reaction IDs (https://www.vmh.life/) (Noronha et al., 2019). h) Healthy microbiomes by body site, i) gut microbiomes of CD patients and healthy controls, j) gut microbiomes of healthy and undernourished children, k) gut microbiomes of PD patients and healthy controls. UMAP = uniform manifold approximation and projection, CD = Crohn’s disease, IBD = inflammatory bowel disease, PD = Parkinson disease, T2D = type 2 diabetes, UC = ulcerative colitis.

When further analysing the generated machine learning classifiers, we found that the relative abundances of at most 20 reactions were sufficient to accurately predict the stratification group for five of the 11 analysed datasets (Figure 4a), while the absolute presence of at most 20 reactions accurately predicted the correct group assignment for five datasets (Figure 4a). For instance, the separation of healthy microbiomes by body site was driven by the relative abundance of only two reactions, i.e., oxalate secretion by diffusion (VMH ID: OXAte) and acetyl-CoA carboxylase (VMH ID: ACCOAC) (Figure 4h). IBD patients and controls could be distinguished by the relative abundance of three reactions, *N*-formylglutamate deformylase (VMH ID: NFORGLUAH), periplasmatic NMN glycohydrolase (VMH ID: NMNt7pp), and acetate proton symport (VMH ID: ACt2r) (Figure 4i). ACt2r, representing the transport of acetate, a key metabolic product of the gut microbiome (Koh et al., 2016), was depleted in Crohn’s Disease (CD) and ulcerative colitis (UC) compared to healthy individuals (Figure 4i). Gut microbiomes of healthy and undernourished children could be distinguished with 97% accuracy by the relative abundances of two reactions, α,α-trehalose-phosphate synthase (VMH ID: TRE6PS) and thymidylate synthase (VMH ID: TMDSf2), both of which were depleted in undernourished individuals (Figure 4j). Gut microbiomes of PD patients and controls could be stratified by a signature of eight reactions (Figure 4k). For the remaining dataset, the classifier retrieved a subset of 16 to 100 reactions whose abundance stratified the groups (Figure S19-S25). When plotting the relative abundance of these reactions, distinct clusters were found for healthy adult and infant gut microbiomes, healthy and premature infant gut microbiomes, and adult gut microbiomes with infection and healthy controls (Figure S19-S21). For the stratification of adult gut microbiomes by country, several clusters were identified, which were each enriched in samples from Denmark, the Netherlands, Mongolia, and Tanzania (Figure S22). In the remaining datasets, less clear clustering was observed (Figure S22-S25), in agreement with the lower accuracy of the classifier for these datasets (Figure 4a). Taken together, the analysis revealed key reactions that could distinguish between microbiomes in health and disease states, body sites, age groups, and countries.

### Model-based statistical analyses further reveal differences between groups

To further elucidate which reactions were driving the separation in these datasets, we performed a statistical analysis for reaction presence, reaction abundance, and subsystem abundance for the 11 datasets (Table S11a-c, Methods). After statistical testing, the p-value was corrected for false discovery rate using the Bonferroni method (Benjamini et al., 2001). For eight of the 11 datasets, more than 10% of total features were significant after false discovery rate correction (Figure 5a). In fact, the relative abundances of >90% of the reactions were significantly different between healthy microbiomes by body site, healthy gut microbiomes by country, and between gut microbiomes of healthy individuals and those with infection (Figure 5a). In contrast, microbiome models of infection patients with and without antibiotics use, of normal weight, overweight, and obesity, and of T2D and healthy patients showed <10% significance of total features. This result is in line with the above-observed less clear separation of groups in these datasets in UMAP clustering (Figure S16-S18) and the lower prediction potential of the classifier for groups in these datasets (Figure 4a). It must be noted that for each dataset, a different total number of features was present due to the different number in individual samples included (Figure 4a, Table S8). We then grouped the enzymatic reactions, for which relative abundance was significantly different across the 11 datasets, by metabolic subsystem (Figure 5b). Metabolic subsystems that made up the biggest contributions to significantly different reactions included xenobiotic metabolism, amino acid metabolism, cell wall biosynthesis, lipid metabolism, carbohydrate metabolism, and vitamin and cofactor metabolism (Figure 5b). A similar distribution by metabolic subsystem was seen for enzymatic reactions that were significantly different in absolute presence across the 11 datasets (Figure S26). Finally, we individually visualised the reactions with statistically significant relative abundances for each dataset (Figure S27-S33). Distinct clusters were observed for microbiomes by body site. Equally, distinct cluster were obtained for gut microbiomes of healthy and premature infants, of healthy and undernourished children, and of healthy adults and those with infection (Figures S27-S30). These results were consistent with the clear stratification of these datasets through UMAP clustering (Figures 4b, d, e, g). For the classification of adult gut microbiomes by country, clusters enriched in samples from Denmark and the Netherlands were observed (Figure S31), in agreement with the independently generated results from the random forests classifier (Figure S22). A less clear separation based on significantly different reactions was observed for gut microbiomes of obese against normal weight individuals and of T2D patients and healthy controls (Figure S32-S33), again in agreement with the random forest classifier (Figures S24, S25). Taken together, for certain comparisons of groups (e.g., disease states) with matched controls, clear differences in metabolic pathway structure and function were found in our microbiome models, while others could be less clearly stratified. As our models take the relative microbial abundances as input, these findings also reflect that microbiome composition was distinct between groups in some datasets (e.g., infection vs. healthy) and less different in others (e.g., T2D vs healthy).

**Figure 5:**
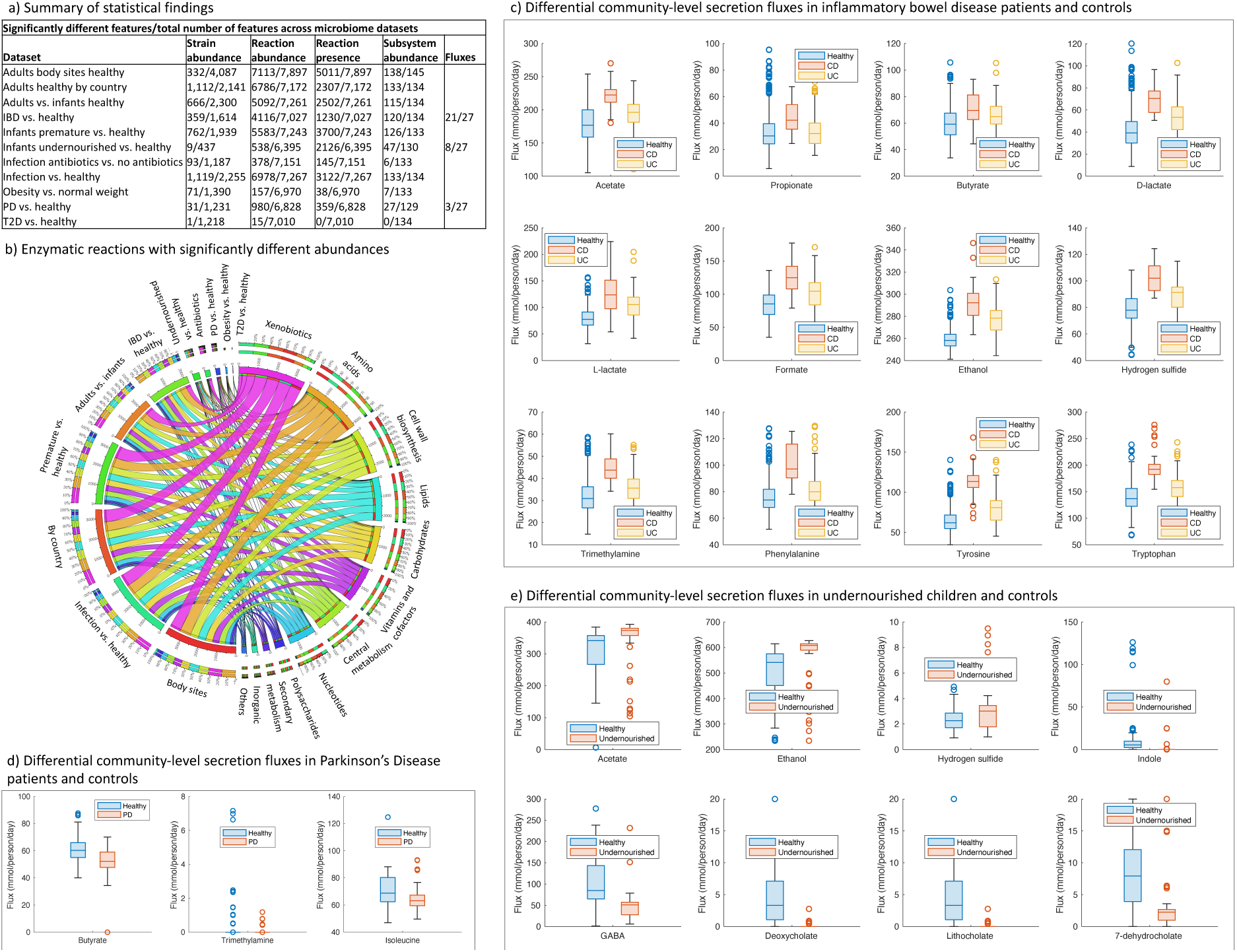
Analysis of personalised microbiome models constructed from APOLLO through statistical analyses. a) Overview of features that were statistically significant out of the total features between groups in the 11 datasets. Details are shown in Table S11a-d. b) Enzymatic reactions that were statistically significant between groups in the 11 datasets by relative abundance shown by corresponding reaction subsystems. c) Community-wide metabolite secretion fluxes that were significantly different between IBD patients and healthy controls. Shown are the 12 metabolites with the highest total production potential. d) Community-wide metabolite secretion fluxes that were significantly different between PD patients and healthy controls. e) Community-wide metabolite secretion fluxes that were significantly different between undernourished children and healthy controls. IBD = inflammatory bowel disease, CD = Crohn’s Disease, UC = ulcerative colitis, PD = Parkinson disease, T2D = type 2 Diabetes.

### Metabolite secretion potential varies between disease states and healthy controls

In addition to determining reaction and pathway abundances, metabolic community modelling also allows for the prediction of the microbiome-wide potential to secrete metabolites of interest, corresponding to faecal secretion (Heinken and Thiele, 2022). Using flux balance analysis (Orth et al., 2010), we retrieved 27 metabolites, which have been found to be altered in previous simulations in IBD (Heinken et al., 2021c) and PD patients (Hertel et al., 2019). Accordingly, we simulated the community-wide production potential for the 27 metabolites in IBD patients and controls, PD patients and controls, and in undernourished children and controls (Methods). Of the 27 computed metabolites, 21, three, and eight were statistically significantly different in their production potential between patients and controls for IBD, PD, and undernourishment, respectively (Figure 5a, Table S11d). The microbiome models of CD patients had an increased production potential for several amino acids, lactate, ethanol, TMAO, and hydrogen sulphide (Figure 5c), in agreement with previous results (Heinken et al., 2021c) and reported findings on altered metabolites in CD (Lavelle and Sokol, 2020; Ni et al., 2017). However, in the present study, the production potential for short-chain fatty acids and secondary bile acids were also higher in CD patients than in healthy controls, in disagreement with previous simulations (Heinken et al., 2021c; Heinken et al., 2019) and general findings on the metabolome in CD (Lavelle and Sokol, 2020). These discrepancies may be due to missing functions in currently unclassified strains in the microbiome models. For PD patients, three metabolites had a reduced secretion potential compared with healthy controls, namely, butyrate, trimethylamine, and isoleucine (Figure 5d). Reduced butyrate levels have been found in faeces of PD patients (Baert et al., 2021; Chen et al., 2022; Unger et al., 2016). For isoleucine, both decreased (Yan et al., 2021) and increased (Vascellari et al., 2020) levels in faeces of PD patients have been reported. The predicted metabolic profiles of undernourished children’s microbiomes showed an increase in acetate and in the cytotoxic metabolites ethanol and hydrogen sulphide, but a decrease in indole, γ-aminobutyric acid, and secondary bile acids compared with healthy control microbiomes (Figure 5e). Reduced faecal amino acid and bile acid levels have been found in protein-starved mice compared with controls (Quinn et al., 2022). Taken together, the biosynthesis potential for health-relevant metabolites were found to be significantly altered in the three compared cohorts of disease-associated and control gut microbiomes.

## Discussion

We present APOLLO, a resource of 247,097 genome-scale reconstructions derived from two large-scale resources of MAGs. APOLLO was built using an established computational pipeline (Figure 1) that systematically explores the metabolic capabilities encoded in metagenome-assembled genomes, allowing for strain- and molecule-resolved analysis of their metabolic potential at unprecedented detail. We demonstrated that taxonomically related strains clustered together by reaction and metabolite content, strengthening confidence in the predictive potential of the generated refined genome-scale reconstructions. We developed a machine learning classifier that could predict taxonomical association from the reconstruction traits and capabilities with high accuracy and that was robust for MAGs from different sources. Based on the classifier, we also identified key reactions stratifying taxa. Finally, to analyse the reconstructed strains in their ecological context, we built and interrogated over 14,000 personalised microbiome models. From these personalised models, we retrieved functional differences in reaction and pathway content between individuals based on health status, body site, geography, and age group. The APOLLO resource is freely available to researchers at https://www.vmh.life/.

Compared with genome-scale reconstructions obtained from annotated reference genomes, such as those from the AGORA2 resource (Heinken et al., 2023), APOLLO presented a lower average percentage of flux consistent reactions (Figure 2a) and a lower metabolic coverage (Figure 2b). With continuing efforts to manually curate MAGs and improve tools for MAG-based reconstruction (Chen et al., 2020; Yang et al., 2021), genome-scale metabolic reconstructions built from these MAGs will also increase in metabolic coverage, accuracy, and flux connectivity. As demonstrated in this work, a clear advantage of the usage of MAGs for reconstructing human microbes is the far more representative taxonomic coverage of the human microbiome compared with reference genome-based resources as up to 50% of human intestinal microbes lack a high-quality reference genome (Yang et al., 2021). Another advantage of MAGs is that they capture individual-specific strain differences and can give rise to host-specific genome-scale reconstructions of microbes and, subsequently, microbiome community models (Frioux et al., 2020). For instance, genome-scale reconstruction based on MAGs could be directly integrated into bioinformatics pipelines to analyse metagenomes (Frioux et al., 2020). A pipeline combining bioinformatics and constraint-based modelling has been already built, which allows for the creation of genome-scale reconstructions directly from metagenomes and the subsequent predictions of interactions between microbes in each sample (Zorrilla et al., 2021). However, it does not cover the semi-automated curation performed by DEMETER (Heinken et al., 2021d). As an alternative, we propose that MAGs could be retrieved from metagenomes through state-of-the-art tools, such as MetaBat 2 (Kang et al., 2015; Kang et al., 2019), subsequently reconstructed using DEMETER (Heinken et al., 2021d), taking advantage of its high-quality, experimental data-guided refinement performed, and analysed as described in this work. It should be noted that the use of manually collected experimental data introduced a curation bias in APOLLO as only 53% of reconstructions could be curated based on available experimental data, compared with 95% in AGORA2 (Heinken et al., 2023), a direct consequence of 64.11% of strains in APOLLO being unclassified on the species level (Figure 2d).

A clear advantage of APOLLO over previous microbiome genome-scale reconstruction resources, such as AGORA2, is that it exhibits relatively better taxonomic distribution of the human microbiome (Lloyd-Price et al., 2016). APOLLO reflects that most human microbial taxa belong to the Bacteroidia and Clostridia classes, and hence, APOLLO allows for a systematic interrogation of the distribution of metabolic traits in these principal taxa. As another advantage, APOLLO also captures currently unclassified genomes, with 7,632 being even unclassified at the phylum level (Table S1). Another significant fraction of reconstructed genomes belongs to currently understudied phyla, such as Verrucomicrobia. For this phylum, only one species, *Akkermansia muciniphila*, has previously been studied in detail. APOLLO contains 1,282 Verrucomicrobia strains unclassified on the species level (Table S1), providing an unprecedented opportunity to study them *in silico* and subsequently, *in vitro*. In future works, interrogating the APOLLO reconstructions may provide important insights into the metabolic capabilities of these understudied and entirely unclassified taxa. Another potential application of APOLLO is the systematic comparison of topological properties, such as metabolite connectivity (Becker et al., 2006) and biosynthetic capabilities across taxa. For instance, AGORA has been previously applied to assess *in silico* vitamin auxotrophies across taxa (Molina Ortiz et al., 2022) and metabolic heterogeneity across strains based on topological features of the reconstructions (Cabbia et al., 2020). APOLLO will enable such applications at a much larger scale and scope.

Previous studies have used metabolic modelling to stratify patients and controls based on individual-specific metabolic fluxes for complex diseases, including PD (Hertel et al., 2019; Rosario et al., 2021), colorectal cancer (Hertel et al., 2021), IBD (Aden et al., 2019; Heinken et al., 2021c), type 1 diabetes (Lamichhane et al., 2022), and cystic fibrosis (Henson et al., 2019). Generally, these studies have involved mapping processed metagenomic sequencing data onto genome-scale reconstructions, such as the AGORA resource (Heinken et al., 2023; Magnusdottir et al., 2017) and constructing personalised microbial community models. In the present study, we have constructed a large resource of personalised microbial community models with an average of 144 mapped strains, capturing over 14,000 samples from all age groups, different body sites, continents, and cases from 21 diseases (Table S7). We provided a systematic analysis of reaction and pathway content across 11 different datasets of microbiome samples matched by disease state, body site, geography, or age group. Numerous microbiome datasets, including from nasal cavity, skin, and vaginal microbiomes, healthy adults and infants, healthy and premature infants, and healthy and undernourished children, showed distinct metabolic traits between the groups, which was confirmed independently by UMAP, the random forest classifiers, and statistical analysis (Figures 4, 5). Other datasets, such as obese and normal weight individuals, and T2D patients and controls, separated moderately to poorly by group (Figures S23-24, S30-31). A reason for the lack of separation may be fewer differences in microbiome abundance between these groups, which consequently translated into fewer differences in metabolic features of the microbiome models that have been parameterised with these relative abundance data (Figure 5a). Through simulations, we delineated the altered metabolic potential of gut microbiomes of CD and PD patients from their respective controls (Figure 5c-e). Our predictions partially confirmed previous metabolomic findings, such as reduced faecal butyrate in PD (Baert et al., 2021; Chen et al., 2022; Unger et al., 2016). The predicted increase in fermentation products and decrease in amino acids and secondary bile acids secretion capabilities of in the gut microbiome models of undernourished children (Figure 5e) represent novel hypotheses, which may be validated in future studies.

Another possible application could be the further contextualisation of the 13,132 gut microbiome models with country-specific dietary regimes. Such simulations may provide mechanistic insights into the known association between Westernised diets and lifestyles, and a loss in gut microbiome compositional and functional diversity (Sonnenburg and Sonnenburg, 2019). As a follow-up, personalised microbial community models could be integrated with whole-body models of human metabolism, which could then be contextualised with dietary information and metadata, including sex, weight, and height (Basile et al., 2022; Thiele et al., 2020). Hence, differences in microbiome structure and function by country of origin, age, and sex could be modelled as well as changes in host-microbiome co-metabolism. The wide variety of captured disease states could also allow for the prediction of potential biomarkers stratifying cases and controls, and therapeutic interventions (Heinken et al., 2021a). Ultimately, our resource of 14,451 microbiome models, due to its unprecedented diversity, could provide novel insights into the interactions between diet, lifestyle, the microbiome, and host health across human populations.

The presented approach used for the generation of APOLLO could be readily applied to other MAG resources beyond the human microbiome. For instance, MAG resources have been published for the microbiomes of wild animals (Levin et al., 2021; Youngblut et al., 2020) and diverse environments on Earth (Nayfach et al., 2021). These genomes could be similarly metabolically reconstructed and modelled, which would elucidate the structure and function of currently largely uncultured microbial ecosystems. Previously, strain-level metabolic modelling could propose experimentally testable growth media for human microbes (Heinken et al., 2014; van der Ark et al., 2018), which may facilitate culturing previously unculturable species. The construction of sample-specific microbiome models would enable the stratification of microbiomes in diverse environments according to their distinct metabolic capabilities. Such systems-level simulations could ultimately lead to novel insights into the adaptation of microbes to their environment and propose manipulations (e.g., through nutrient availability) that could favourably influence the metabolic capabilities of globally important ecosystems, such as the human gut, rumen, ocean, or soil.

## Author contributions

IT, AH, and CT designed the study. AH, AOB, NFOS, CG, EM, FMD, IL, MB, PE, RD, RF, RMD, SH, SRMA, TK, FM, CT, and IT built the draft reconstructions. AH, AOB, CG, EM, FMD, IL, MB, NFOS, PE, RD, RF, RMD, SH, SRMA, and TK collected input data. AH built and tested the refined reconstructions. AB performed taxonomic profiling of metagenomic data. AH built the microbiome models and performed simulations. AH, TOH, BN, FM, and IT analysed data. TOH performed machine learning analyses. BN and FM performed statistical analyses. AH, TOH, BN, and AB drafted the manuscript. AH, TOH, AB, CCT, and IT revised the manuscript. IT, AH, CCT, and R.M.T.F. supervised the study. IT acquired funding.

## Methods

All simulations were carried out in MATLAB version 2021b (Mathworks, Inc.) using IBM CPLEX (IBM) as the linear programming solver.

### Retrieval of reconstructed genomes

The metagenome-assembled genomes (MAGs) used in this study had been published previously (Almeida et al., 2019; Pasolli et al., 2019). Download links for 154,723 MAGs (Pasolli et al., 2019) (“Pasolli” resource) as well as taxonomic assignments for the same genomes were retrieved from https://opendata.lifebit.ai/table/SGB. Another resource of 92,143 MAGs (Almeida et al., 2019) (“Almeida” resource) as well as taxonomic assignments for the same genomes were downloaded from http://ftp.ebi.ac.uk/pub/databases/metagenomics/umgs_analyses/. A further 553 genomes were contained within the Human Gastrointestinal Bacteria Culture Collection (Forster et al., 2019), of which 327 had already been reconstructed as part of the AGORA2 (Heinken et al., 2023) resource. The remaining 226 genomes were downloaded from http://ftp.ebi.ac.uk/pub/databases/metagenomics/umgs_analyses/. Taken together, this resulted in 247,092 genomes, for which genome-scale reconstructions were created as described below.

### Generation of draft reconstructions

Draft reconstructions were created through the KBase (Arkin et al., 2018) online platform in a joint effort by utilising the Narrative interface. To efficiently distribute the effort between all team members, the list of all genomes to reconstruct was divided into batches of 500 to 1,500 genomes through custom MATLAB scripts. For each batch of genomes, draft reconstructions were created by the respective team members through a dedicated Narrative instance. To facilitate this process, a template Narrative (available at kbase.us under https://narrative.kbase.us/narrative/82195) was created and distributed to all team members and consisted of the following steps: (i) download of the assemblies in FASTA format to the Staging Area. For the Pasolli resource, this was performed via the Upload File to Staging from Web - v1.0.12 app after retrieving download links from https://opendata.lifebit.ai/table/SGB. For the Almeida resource, genomes were downloaded from http://ftp.ebi.ac.uk/pub/databases/metagenomics/umgs_analyses/, imported in zipped format into the Staging Area, and unpacked through the Unpack a Compressed File in Staging Area - v1.0.12 app. (ii) Import of the Assemblies into the Narrative through the Batch Import Assembly from Staging Area app. (iii) Annotation of the imported assemblies through the Annotate Multiple Microbial Assemblies app with the Assembly Set created in the previous step as the input. (iv) Creation of draft metabolic reconstructions from the annotated genomes through the Build Multiple Metabolic Models app with the Genome Set created in the previous step as the input. (v) Export of the created draft reconstructions through the Bulk Download Modelling Objects app. To streamline the efforts, all team members reported their progress through dedicated spreadsheets. If a batch could not be processed completely, unreconstructed genomes were retrieved and redistributed into new batches through custom MATLAB scripts.

### Refinement of draft reconstructions

All created draft reconstructions were subsequently refined through the DEMETER (Heinken et al., 2021d) pipeline. Briefly, DEMETER is a COBRA Toolbox (Heirendt et al., 2019) extension that ensures that the resulting refined genome-scale reconstructions conform to the quality standards in the field (Lieven et al., 2020; Thiele and Palsson, 2010a), e.g., correct reconstruction structure, uniform namespace, and thermodynamic consistency, and capture the known biochemical and physiological traits of the target organisms (Heinken et al., 2021d). Moreover, through quality control steps, thermodynamic feasibility, appropriate biomass composition, and correct reconstruction structure are ensured (Heinken et al., 2021d). The amount of curation against experimental data that could be performed for each strain depended on the taxonomic assignment performed by the original authors (Almeida et al., 2019; Pasolli et al., 2019) with unclassified and unnamed species only allowing for limited curation of the respective genome-scale reconstructions. Of the 568 and 116 named species included in the Pasolli and Almeida, respectively, 99 and three, respectively, were not yet accounted for in the input data for DEMETER. Hence, experimental data, namely carbon sources, fermentation products, growth requirements, and consumed and secreted metabolites, were manually collected for these species where available as described previously (Heinken et al., 2023). For genomes that were only classified at the genus level, curation against experimental data was only performed for fermentation products, for which data was propagated from strains of the same genus if available. Genomes classified only at the family level or higher could not be curated against experimental data. However, their biomass composition was curated based on gram status and periplasmic transport reactions were added accordingly (Heinken et al., 2023; Heinken et al., 2021d). After passing through the refinement pipeline, the reconstructions were extensively tested for agreement with available experimental data, biomass production on a complex medium, and reasonable ATP production on the complex medium as described previously (Heinken et al., 2023). Functions in DEMETER were continuously revised to account for the new reconstructions. Overall, the pipeline resulted in 247,092 semi-automatically refined genome-scale reconstructions.

### Computation of strain-specific metabolic features

For all 247,092 genome-scale reconstructions, metabolic properties of the corresponding models were computed using functions implemented in DEMETER (Heinken et al., 2021d) and in the COBRA Toolbox (Heirendt et al., 2019). Specifically, the distribution of metabolites and reactions across models was computed through the getReactionMetabolitePresence function. The function reports the distribution of all unique reactions in a reconstruction resource for each model with “1” indicating presence and “0” indicating absence. The theoretical metabolite uptake and secretion potential on unlimited medium, consisting of every metabolite the microbe could potentially transport, were computed through the computeUptakeSecretion function. Briefly, the function computes the minimal and maximal fluxes for each exchange reaction in a model through flux variability analysis (Gudmundsson and Thiele, 2010). Metabolites are considered consumed if the minimal flux through the corresponding exchange reaction is negative and considered secreted if the maximal flux through the corresponding exchange reaction is positive. Note that metabolites can be both consumed and secreted by the same model. Finally, the potential to internally synthesise metabolites on unlimited medium regardless of transport capability was computed for all cytosolic metabolites through the computeInternalMetaboliteProduction function. The function creates a demand reaction for each metabolite in a model and maximises its flux through flux balance analysis (Orth et al., 2010). Metabolites with a positive flux through the corresponding demand reaction are considered internally produced. For both uptake and secretion potential and internal production potential, we used qualitative data for further analyses, with −1 representing uptake, 1 indicating secretion or production, and 0 indicating neither.

### Development of the machine learning classifier

To build a classifier that could predict strain taxonomy based on the strain-specific metabolic features described above, a Random Forest model was chosen to classify the data. The data was divided with an 80:20 split into training and validation groups. The groups were stratified to ensure similar levels of each taxonomic assignment would be present amongst both groups.

Hence, taxonomic assignments with few instances were always present in the validation group and that there was no over-abundance of taxonomic assignments with many instances. Gini impurity was used as the feature importance metric. Potential feature redundancy was determined by calculating the Spearman’s rank-order correlation for the features. All but one of the closely related features can be removed with minimal impact to accuracy, thus, reducing the dataset size further. To visualise the separation between strain, linear discriminant analysis (LDA) was used to find an optimum representation of the reduced dataset in three-dimensional space. Uniform manifold approximation and projection for dimension reduction (UMAP) (McInnes et al., 2018) was used to visually show that the dataset has inherent divisibility. The classifier was built in Python version 3.8.0 and relied on the pandas, fastai, sklearn, matplotlib and scipymodules.

### Taxonomic profiling

Taxonomic profiling was performed for a subset of MAGs from the Pasolli resource that had been retrieved from nasal cavity, skin, and vaginal microbiome metagenomes samples originally part of the Human Microbiome Project (HMP) (Human Microbiome Project, 2012). Abundances of the HMP samples were retrieved. We then divided the HMP samples and the MAGs according to their body origins. For each body origin, a dedicated ad-hoc database was created with the Bowtie2-build (v 2.4.2) (Langmead and Salzberg, 2012) function. Each database comprises the MAGs corresponding to the HMP samples for each body source and the genomic sequences used for the AGORA2 (Heinken et al., 2023) collection. The raw reads were mapped on the appropriate database using Bowtie2 −q (v. 2.4.2) (Langmead and Salzberg, 2012). The SAM files of the alignment were subsequently converted to BAMs, ordered BAMs and indexed BAMs through the SAMtools (v 0.1.19) (Li et al., 2009) set of utilities. Finally, the coverage of each scaffold and therefore the abundance of each MAG in the samples under examination was retrieved with the CheckM genome suite (v. 1.1.3) (Parks et al., 2015) using the functions CheckM coverage and CheckM profile.

### Construction and interrogation of personalised microbiome models

Relative strain-level abundances that had been determined in 13,332 gut microbiome metagenomic samples for 1,952 MAGs included in the Almeida resource (UMGS genomes) and 553 human gut reference (HGR) genomes (Almeida et al., 2019) were retrieved from http://ftp.ebi.ac.uk/pub/databases/metagenomics/umgs_analyses/mapping_results/bwa_cover <u>age.csv</u>. Of these 2,505 genomes, genome-scale reconstructions were created for 2,178 in the present study, and 327 had been reconstructed as part of AGORA2 (Heinken et al., 2023). Genomes from the retrieved abundance data were mapped onto the nomenclature of the corresponding genome-scale reconstructions, and strains with abundances below 0.001 were removed in the respective samples. Through the mgPipe workflow implemented in the Microbiome Modelling Toolbox 2.0 (Heinken and Thiele, 2022), personalised microbiome models were generated for 13,332 gut microbiomes. Moreover, personalised microbiome models were generated for 87 nasal cavity, 1,143 skin, and 89 vaginal metagenomic samples based on the taxonomic profiling performed in the present study for a subset of 1,021 MAGs in the Pasolli resource and 1,046 reference genomes included in AGORA2. Relative reaction abundances and absolute presence of all unique reactions present in at least one of the 14,451 microbiome models, as well as relative abundances on the subsystem level, were computed through the mgPipe workflow. We then defined a set of metabolites of interest that were known to be microbial in origin and. had been previously linked to disease states (Table S9). For a subset of 916 microbiome models, secretion capabilities of the 27 metabolites by each microbial community model were predicted using the analyseObjectiveShadowPrices function in the Microbiome Modelling Toolbox 2.0.

### Definition of microbiome datasets

To compare results obtained from the 13,332 created gut microbiome models in the context of the corresponding host’s age, country of origin, disease status, and other available host information, metadata provided by Almeida et al. for the was retrieved from https://static-content.springer.com/esm/art%3A10.1038%2Fs41586-019-0965-1/MediaObjects/41586_2019_965_MOESM3_ESM.xlsx. To obtain more information on each sample, the GMRepo database (Dai et al., 2022) was searched using NCBI Project IDs provided by Almeida et al. Where possible, the original publications were searched for additional information on the samples. Table S7a shows the collected metadata for all samples. Based on the collected metadata, we defined 11 datasets within the gut microbiome samples as follows. We first retrieved cohorts of patients and matched controls within the 13,332 gut microbiome samples for categories of diseases that had been modelled previously (Aden et al., 2019; Baldini et al., 2020; Heinken et al., 2021c; Heinken et al., 2019; Hertel et al., 2019; Lamichhane et al., 2022; Rosario et al., 2021; Shoaie et al., 2015), namely, CD, obesity, T2D, and PD (Figure 4a, Methods). To model microbiomes by body site, we defined a comparison by body site by including 87 nasal cavity, 1,143 skin, and 89 vaginal from healthy adults. To evaluate the impact of host age and infant developmental status, we compared (i) all known healthy adult and infant gut microbiomes, (ii) all healthy and premature infant gut microbiomes, and (iii) gut microbiomes from a cohort of malnourished and control infants. To determine if differences in composition translate into functional differences between Westernised and non-Westernised microbiomes, all healthy adult gut samples were stratified by country of origin. To predict the effect of infection and antibiotic use on gut microbiome structure and function, we compared the gut microbiomes of infection patients with and without antibiotics use as well as healthy controls.

Where possible, samples were matched to their respective controls from the original studies that had generated the metagenomes, using NCBI Project IDs as reference. In cases where the original studies did not contain matching controls, multiple studies were pooled. This was the case for healthy adult and infant gut microbiomes, premature and healthy infant gut microbiomes, healthy adult gut microbiomes by country, and healthy and infection gut microbiomes. Samples with unknown age, country of origin, or health status were not considered. It was possible to pool samples from multiple studies as all metagenomes had been processed through the same workflow. It was confirmed that the compared groups in the datasets consisting of pooled samples were distinct in their reaction content. Table S7b shows the samples belonging to each of the 11 defined datasets.

### Statistical analysis

To find statistically significant results for the eleven subgroups, the means and standard deviations were taken for the total metagenomic read counts, α-diversity, β-diversity, Shannon diversity, Simpson diversity index, total number of reactions and metabolites, reaction presence, microbial, subsystem and reaction relative abundance and if available simulated microbiome excretion fluxes for a subset of 27 reactions. For reaction presence Fisher’s exact test and the Chi-square test were used to calculate significance. If a reaction was present in more than 95% or less than 5% of all samples in a subgroup, the reaction was excluded from analysis. For the other parameters a student t-test or ANOVA was used depending on the amount of stratification groups. If the standard deviation for the relative abundance of subsystems or reactions was equal to 0, they were excluded from analysis. All p-values were then corrected by the Bonferroni method and deemed significant if the corrected p-value was below 0.05.

## Data visualisation

Venn diagrams were generated with the Venn Diagrams online tool (https://bioinformatics.psb.ugent.be/webtools/Venn/). Circle plots were generated through the Circos online tool (http://mkweb.bcgsc.ca/tableviewer/visualize/). UMAP and LDA plots were generated through the Python modules umap-learn (https://umap-learn.readthedocs.io/en/latest/) and sklearn (https://scikit-learn.org/stable/). All other data was visualised in MATLAB version 2021b (Mathworks, Inc.) and in R version 4.1.0 (Team, 2013) using R Studio (Team, 2020).

## Data and Software Availability

All scripts created for this study are available at https://github.com/ThieleLab/CodeBase. The APOLLO reconstructions and the 14,451 personalised microbiome models are available at https://www.vmh.life/ (Noronha et al., 2019) (upon publication).

